# DNAlogo: a smart mini application for generating DNA sequence logos

**DOI:** 10.1101/096933

**Authors:** Yabin Guo

## Abstract

Sequence logos are frequently used for presenting consensus sequences and motifs of nucleic acids and proteins. WebLogo of UC Berkeley is the most popular tool for creating sequence logos, whereas, it is often restricted by the internet speed in some developing countries and no graphic interface for its stand-alone version. Here, the author generated an application, DNAlogo, using VB.net, which runs in Windows system. DNAlogo is small and convenient. It creates both bitmap and vector map. Beside the classic sequence logo function, DNAlogo introduced compensated logo to generate consensus sequences considering different GC contents of different genomes. DNAlogo provides a simple way for researchers without programming knowledge to create DNA sequence logos.

---

Sequence logo is a powerful tool for presenting consensus sequences or motifs of nucleic acids and proteins (Schneider and Stephens 1990). WebLogo, a web-based sequence logo generator hosted by the University of California, Berkeley is the most popular logo generator so far (Crooks *et al.* 2004). WebLogo has a graphical interface and is convenient and highly configurable. However, its application is occasionally restricted by the internet speed, especially in developing countries. Moreover, when the sequence number exceeds 10,000, a command line interface will have to be used instead of graphical interface, but many users in biological sciences fields found it difficult to perform the installation and configuration of WebLogo and GhostScript (for vector map output) due to lacking relative knowledge. Here I made an application, DNAlogo, which creates DNA sequence logos in Windows with a graphical interface. DNAlogo is written in VB.net based on the algorithm and correction described previously (Schneider *et al.* 1986; Schneider and Stephens 1990).

The input for DNAlogo can be either Fasta format or pure sequences (sequence per line). DNAlogo can also read matrices designating the bit values of each character directly. The bitmaps showed in the picture box of DNAlogo can be saved as JPEG format. For publication purpose, vector maps can be output by saving PostScript (.ps) files and further processed using Adobe Illustrator.

Similar to WebLogo, DNA logo can create both bit logos and frequency logos. Frequency logos contain only the information of frequencies of the four nucleotides, but no information of sequence conservation.

Because GC contents vary among genomes, DNAlogo added a new function to minimize the bias introduced by different GC contents, which is named “Compensated logo”. For example, in fission yeast *Schizosaccharomyces pombe* genome, the GC content is only 36%, in which even random sequences show an AT rich pattern in sequence logos. However, random sequences should have no conservation. Fig. 1 showed the sequence pattern of the previously reported integration sites of Tf1 transposon in *S. pombe* (Guo and Levin 2010). It seems A or T is preferred in all of the positions when presented using a regular logo (Fig. 1A), but when the logo is compensated by the GC content (36%), preferences for G or C were showed in certain positions (Fig. 1B). To my knowledge, this is the first time that the GC contents were considered in sequence logo generation.

**Fig. 1.**
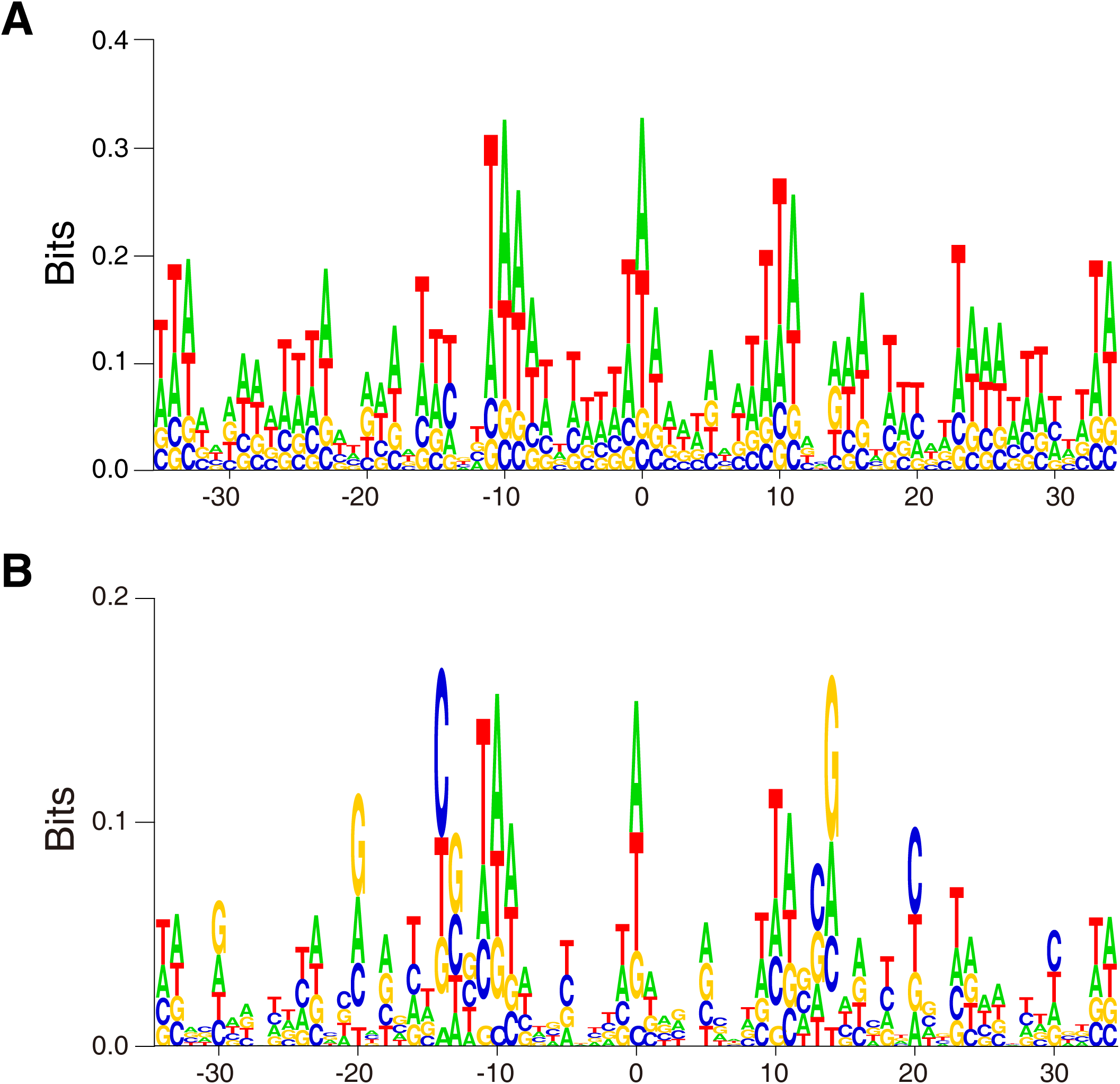
50,613 *S. pombe* genomic sequences flanking Tf1 integration site were aligned according to the Tf1 target site duplications and orientations, and sequence logos were created using DNAlogo. A, Regular logo; B, Compen-sated logo.

Actually, some sequence logos in two previous publications were created using DNAlogo (Guo *et al.* 2013; Chatterjee *et al.* 2014). Logos in Figure 8 of the latter are compensated logos.

DNAlogo is small, convenient, and no installation is needed. People can use DNAlogo without any programming or bioinformatics knowledge. The DNAlogo application (.exe file) and user’s manual were deposited in Github (<u>https://github.com/DNAworker/DNAlogo).</u>

